# Different types of social links contrastingly shape reproductive wellbeing in a multi-level society of wild songbirds

**DOI:** 10.1101/2024.02.02.578606

**Authors:** Samin Gokcekus, Josh A. Firth, Ella F. Cole, Ben C. Sheldon, Gregory F. Albery

## Abstract

The social environment has diverse consequences for individuals’ welfare, health, reproductive success, and survival. This environment consists of different kinds of dyadic bonds that exist at different levels; in many social species, smaller social units come together in larger groups, creating multilevel societies. In great tits (*Parus major*), individuals have four major types of dyadic bonds: pair mates, breeding neighbours, flockmates, and spatial associates, all of which have been previously linked to fitness outcomes. Here, we show that these different types of dyadic bonds are differentially linked with subsequent reproductive success metrics in this wild population and that considering spatial effects provides further insights into these relationships. We provide evidence that more social individuals had a higher number of fledglings, and individuals with more spatial associates had smaller clutch sizes. We also show individuals with stronger bonds with their pair mate had earlier lay dates. Our study highlights the importance of considering different types of dyadic relationships when investigating the relationship between wellbeing and sociality, and the need for future work aimed at experimentally testing these relationships, particularly in spatially structured populations.

## Introduction

Many animals live in complex societies (Ward & Webster 2016). The social behavior of individuals within populations gives rise to structured social systems, and social interactions influence individuals’ decisions and behaviors, including foraging activity, mate choice, survival, and reproduction (Snyder-Mackler *et al*. 2020). Consequently, variation in sociality is an important determinant of overall fitness (Bond *et al*. 2021; Ellis *et al*. 2019; Menz *et al*. 2020; Siracusa *et al*. 2020; Turner *et al*. 2020). While social relationships come in many forms, sociality is often quantified using a single type of dyadic bond between individuals, for example by quantifying the strength of bonds between reproductive pairs, foraging partners, or group members. A key remaining question is how individual variation in different types of dyadic relationships simultaneously shape fitness outcomes in these complex societies.

Different kinds of dyadic relationships within populations can pave the way for hierarchically nested social relationships that form multilevel societies (Papageorgiou & Farine 2021). Indeed, all representations of social relationships are ultimately built upon dyadic units. Multi-tiered social structures often consist of breeding pairs or families that are nested within higher-order groups, allowing for flexibility in social interactions (Grueter *et al*. 2017). Smaller, more stable units may come together to form larger fluctuating groups, such that individuals may maintain and keep track of relationships at multiple levels (Papageorgiou *et al*. 2019). Assessing how these layers influence fitness is crucial to gaining a more complete understanding of understanding how individual-level selection on behaviours build up to produce societal structure. However, quantifying these effects in wild populations can be difficult, as it requires fine-scale behavioural data from multiple social contexts across years alongside fitness outputs. Long-term, individual-based studies with high-resolution data are particularly useful in this regard (Clutton-Brock & Sheldon 2010; Sheldon *et al*. 2022).

The long-term study system of great tits (*Parus major*) at Wytham Woods provides an ideal opportunity to link different kinds of social relationships with fitness outcomes. Individuals are monitored during winter and spring over multiple years and in multiple social contexts, with associated data collected on reproductive success. As detailed below, four broad types of associations exist within this population that are known to be biologically meaningful: pair mates (Culina *et al*. 2015b, a, 2020; Firth *et al*. 2018a; Firth & Sheldon 2016), breeding neighbours (Gokcekus *et al*. 2023; Grabowska-Zhang *et al*. 2012a, b), winter flockmates (Firth *et al*. 2015, 2017, 2018b), and winter spatial associates (Farine *et al*. 2015; Firth *et al*. 2018b; Firth & Sheldon 2016).

In this monogamous species, **pair mates** bond and defend their territory during the breeding season, while also staying close during the non-breeding season (Firth *et al*. 2015). Pair bonds can be understood as the core unit of this multilevel society (Grueter *et al*. 2020). Spatial and social associations often have an influence on mate choice (Beck *et al*. 2020), and mate choice will in turn have an influence on fitness (Fayet *et al*. 2017; Griffith *et al*. 2011; Ihle *et al*. 2015). Stronger associations within the pair bond, often assessed in terms of familiarity over time, seem to come with fitness benefits in many species (Ausband 2019; Van De Pol *et al*. 2006; Sánchez-Macouzet *et al*. 2014). In this population, pairs who meet earlier in the winter have higher reproductive success (Culina *et al*. 2020), pairs that stay together longer are more likely to survive (Culina *et al*. 2015a), and individuals that are unsuccessful are more likely to divorce (Culina *et al*. 2015b; Gokcekus *et al*. 2023).

During the breeding season, territorial great tits have **breeding neighbours,** and these neighbours often also coexist in winter flocks (Firth & Sheldon 2016). Studies have shown that familiarity among neighbours can lead to mutual tolerance and decreases in aggression and energy required for territorial defence, potentially optimizing the timing of reproduction and facilitating coordination and cooperation (Bebbington et al. 2017; Beletsky & Orians 19s89; Booksmythe et al. 2012; Grabowska-Zhang et al. 2012a; Liebgold & Cabe 2008; Riehl & Strong 2018; Siracusa et al. 2020) (but see Müller & Manser 2007; Temeles 1994). Great tits in Wytham Woods benefit from having familiar neighbours (individuals that were neighbours in previous years), through increases in cooperation (Grabowska-Zhang *et al*. 2012a) and, presumably consequentially, higher reproductive success (Grabowska-Zhang *et al*. 2012c).

Outside of the breeding season, individuals also have **flock mates** that they forage with in relatively large fission-fusion winter flocks (Massen *et al*. 2010). Studies in mammalian and avian species have demonstrated that having more (Cameron *et al*. 2009; Ellis *et al*. 2019; McFarland *et al*. 2017; Turner *et al*. 2020) or stronger (Cheney *et al*. 2016; Kohn 2017; Silk *et al*. 2010) social associations can come with fitness benefits (but see Menz *et al*. 2020; Sabol *et al*. 2020). It is possible that more/stronger bonds allow for more appropriate individual or group responses to social and environmental stressors (Campbell *et al*. 2018; Micheletta *et al*. 2012; Young *et al*. 2014), higher rates of cooperation (Berghänel *et al*. 2011; Samuni *et al*. 2018, 2021), and/or increases in dominance that lead to more successful reproduction (Bray *et al*. 2021; Feldblum *et al*. 2021; Schülke *et al*. 2010; Strauss & Holekamp 2019). In this system, an individual’s flock mates influence patterns of social information acquisition during winter foraging (Firth *et al*. 2016).

During the winter, individuals are also **spatially associated** with others while foraging, which may indicate some form of association (or non-independence) but not necessarily direct social interactions. The effects of such associations can vary based on factors such as predation risk and competition (Alberts 2019). The configuration and constraints of the external environment can also influence spatial proximity, and therefore associations, between individuals (He *et al*. 2019; Papageorgiou *et al*. 2019). Partitioning out these purely spatial associations can help to gain a clearer understanding of how different dyadic relationships contribute to the formation of multi-level societies and their fitness consequences.

Finally, all of these links occur within the context of the **spatial environment**, which plays a role in “bottom-up” determination of the social network itself (Albery *et al*. 2021; Webber *et al*. 2023). Because the social network’s formation can be confounded with a variety of environmental (Fisher *et al*. 2021) and demographic drivers (Shizuka & Johnson 2019), many of which can also influence fitness, it is important to take the spatial environment – and particularly population density – into account when understanding sociality-fitness links.

Although the link between sociality and fitness has been considered for several types of dyadic relationships separately, considering them together and in concert with spatial effects is necessary to gain a better understanding of how natural selection operates on individual social behaviour, how this varies for different types of relationships, and therefore how evolution shapes social organization. What are the consequences of different types of sociality? Here, we examine how these four types of relationships contribute to multiple components of fitness within a multi-level society of wild songbirds, while considering spatial effects based on breeding position.

## Methods

### Study system

Our study was conducted in Wytham Woods, Oxford, UK (51°46’N, 1°20’W) on the long-term study population of great tits (*Parus major*). These 385ha of mixed deciduous woodland house ∼1018 fixed nest boxes, with known GPS coordinates, that are monitored yearly to record breeding attempts and performance (Perrins 1965). Either during the breeding season (at the nest) or throughout the year (through mist netting) individuals are caught, fitted with a unique BTO (British Trust for Ornithology) metal leg ring and a plastic leg ring with a passive integrated transponder (PIT) tag, aged and sexed, and measured to record standard morphometric information. It is estimated that 90% of the population is tagged (Aplin *et al*. 2013).

### Data collection

#### Winter social network data collection

During the winter, great tits feed from patches of resources that are distributed throughout the woodland. Sunflower seed feeding stations with two opposing access holes equipped with RFID antennae (Dorset ID, Aalten, The Netherlands) were placed in a stratified grid at 65 fixed locations approximately 250m apart. Feeders opened automatically every weekend over the winter (December-February, 2011-2014) from pre-dawn to after dusk. When birds used one of these feeders, tags allowed the detection of the time and date of their visit, creating a spatio-temporal data stream consisting of each individual visit to a feeder.

### Breeding season data collection

#### Fitness measures

During the breeding season (March-June), birds breeding in nest boxes were monitored through the stages of nest building, egg laying, incubation, and rearing offspring. Nests were checked weekly until eggs were found. Based on the assumption that one egg is laid each day, lay date was recorded (if one egg was found) or estimated (if more than one eggs were found) as the date the first egg of the clutch was laid. After incubation began, clutch size was recorded as the maximum number of eggs within the nest. Parents were either identified or ringed a week after eggs hatched, and all chicks were ringed and weighed a week later.

Subsequently, nests were checked to determine the number of chicks that had fledged in each breeding attempt.

#### Labelling neighbours

In order to label territorial neighbours during the spring, we used the spatial arrangement of occupied nest boxes to estimate individual territories and their boundaries. For each occupied box, a Voronoi diagram (Thiessen polygon) was drawn to include all of the points that were closer to the focal box than any other occupied box (Adams 2001; Schlicht *et al*. 2014). This method of estimating territories and neighbours accurately accounts for population density and is highly correlated with territory sizes and boundaries that are manually determined (Grabowska-Zhang *et al*. 2012c; Schlicht *et al*. 2014; Wilkin *et al*. 2006) as well as with the winter social structure (Farine & Sheldon 2015; Firth & Sheldon 2016).

#### Calculating the four measures of dyadic associations

Each of the four measures of dyadic bonds was calculated based on the social interactions of individuals during the winter. In the spatio-temporal data stream created by the feeder visits, intermittent bursts of clustered activity can be detected which denote flocks that are arriving to feed together. These flocks were detected using a Gaussian mixture model (GMM) that statistically assigns each individual visit to the group that it is most likely to belong to (Psorakis *et al*. 2012). We created a social network (association matrix) for each winter based on co-occurrences in flocks using the simple ratio index (SRI). The SRI values range from 0 to 1 and can be defined by the following equation: SAB = x/(x + yAB + yA + yB), where:

- SAB is the strength of association between A and B
- x is the number of times A and B were in the same flock
- yAB is the number of times A and B were detected at the same time but not together
- yA is the number of times A was detected in a flock without B
- yB is the number of times B was detected in a flock without A

To quantify the relationship between **pair mates**, we only retained records where the identity of both individuals in the breeding pair was known. Based on the winter social network, we calculated the strength of each pair’s bond during the winter prior to the breeding season (following Culina *et al*. 2020; Firth *et al*. 2018a).

The relationship between **breeding neighbours** was quantified based on the territory and neighbour estimates described above and winter social networks. For every focal and each of their first-order neighbours, we calculated the bond strength between them from the previous winter (following Firth & Sheldon 2016). For each individual, we calculated 3 metrics: the average bond strength to their male neighbours, female neighbours, and all neighbours (male and female combined).

To represent **flockmate** associations, we used two social network metrics, weighted degree (the number of connections an individual had weighted by the strength of those connections i.e. the sum of their SRI scores to others) and average bond strength (the average strength of connections i.e. the mean of their non-zero SRI scores to others). In other words, these metrics are based on co-occurrence within the foraging groups identified through the GMMs described above.

To estimate winter **spatial associations**, we calculated the spatial temporal overlap for every possible dyad by accounting for the number of times individuals were observed at each feeder regardless of the specific foraging flocks they occurred in whilst at the feeder (following Firth *et al*. 2018a). So, for example, if two individuals spent equal proportions of time in the same location on the same day (not necessarily at the same time), they would be assigned a 1; if they were never at the same location on the same day, they would be assigned a 0. Proportions for each focal individual were summed to create a single variable accounting for both the number of individuals each focal was in the same location as and the amount of time they each spent there. In contrast to the flockmate associations, this measure is based only on spatial temporal overlap in location.

To examine the role of socio-spatial demographic distributions, we additionally calculated local **density**. Using the adehabitathr package in R, we created annual space use distribution kernels for the population based on their breeding locations and assigned each individual a local density measure based on the individual’s location on that distribution. This follows a protocol established for badgers and deer (Albery *et al*. 2020, 2021).

### Data analysis

We analyzed three years (2011-2014) of data from 754 individuals. We ran separate models for males and females for each of the five fitness variables derived from the breeding data (binary success, clutch size, lay date, mean chick weight, and number of fledglings), as each individual has different social characteristics (following Gokcekus *et al*. 2023). We fitted Generalized Linear Mixed Models (GLMMs) using the Integrated Nested Laplace Approximation (INLA) R package and fitted the SPDE random effect (for breeding season locations). This models similarity emerging from the distance between points to account for spatial autocorrelation in the response variable (Albery *et al*. 2019; Rue *et al*. 2009); we used locations of nest boxes. Continuous variables were scaled to have a mean of 0 and a standard deviation of 1. The base model included year (as a categorical fixed effect), categorical age (juvenile vs. adult), and habitat quality (the number of oak trees within a 75m radius of the nest box). We also included individual identity as a random effect.

To investigate the effect of each of the dyadic relationship measures, we iteratively added the social effects (one measure of the social pair bond, three measures of breeding neighbour bonds, two measures of flockmate bonds, and three measures of spatial associations; correlation matrix in Supplementary Figure 1) the base model, and used Deviance Information Criterion (DIC) to identify the best fit. In each round, social effects were individually added until all had been included or their addition did not improve the model, using a cutoff of 2 DIC (following Albery *et al*. 2022; Gokcekus *et al*. 2023). Fixed effect estimates were provided by the mean and 95% credibility intervals of the posterior estimate distribution. Significance was determined by examining each effect’s overlap of the 2.5% and 97.5% posterior estimates with zero.

## Results

We analysed data on 754 individuals embedded within 377 mated pairs with a total of 4,498 neighbour bonds and flockmate associations for each individual determined from 204,838 winter foraging flocking events. Our measures of sociality (in terms of the four types of dyadic bonds) influenced three of the five fitness-related variables (i.e. were retained by the model selection process for these models; lay date, clutch size, and number of fledglings).

### Four levels of dyadic bonds

Individuals with stronger winter bonds to their pair mate had earlier lay dates, and this effect remained significant when controlling for breeding spatial autocorrelation (Figure 1; Female: -.13, CI: -.19, -.07; Male: -.13, CI: -.19, -.07).

**Figure 1.**
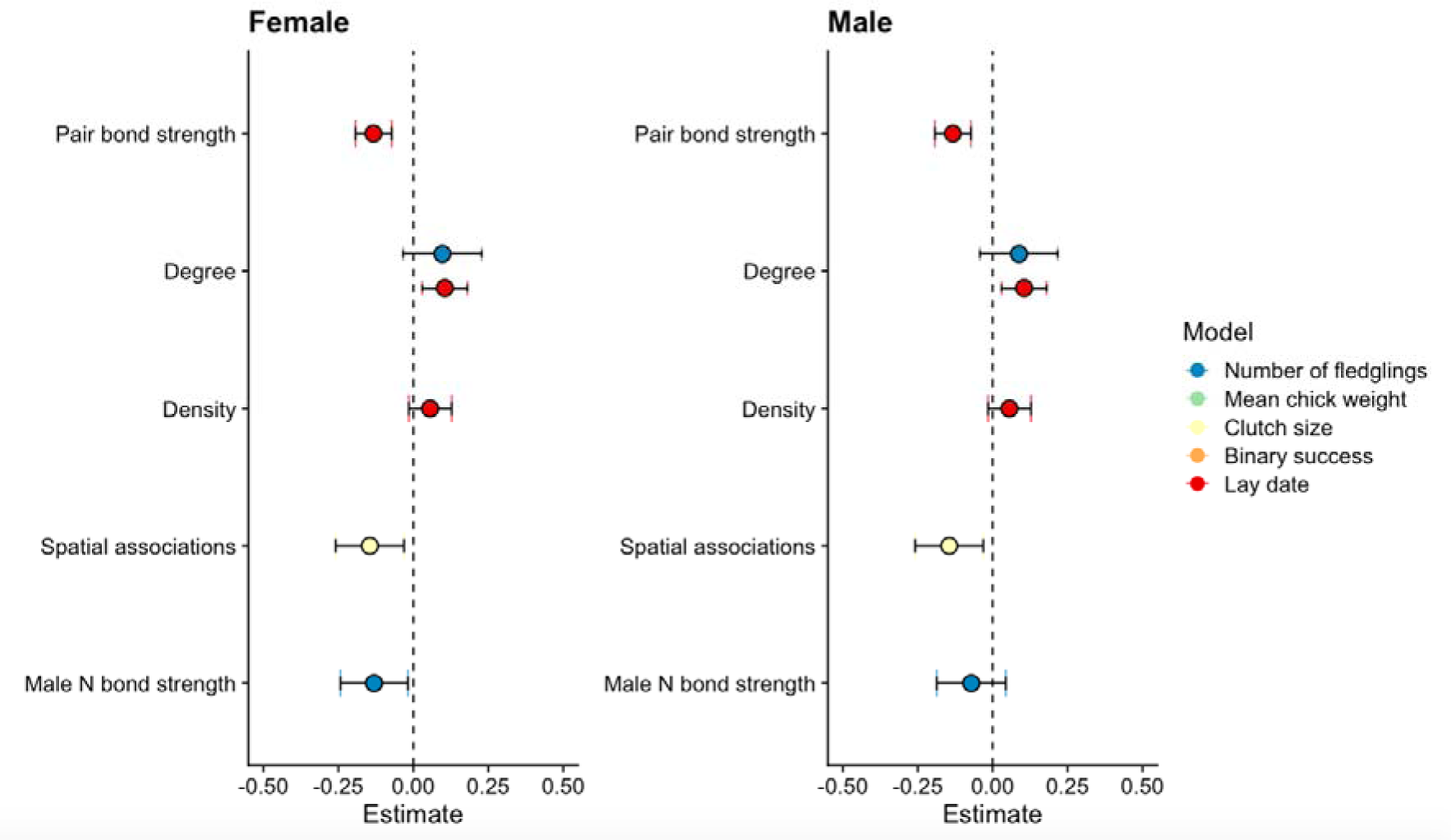
– Summary of the social effects for all ten models, for females and males, with an SPDE effect to account for spatial autocorrelation. Points represent the estimate for each effect that was retained in the model selection process; error bars denote 95% credibility intervals.

Individuals with stronger bonds to their male breeding neighbours had fewer fledglings. This effect was not significant after controlling for breeding spatial autocorrelation for males (-.08, CI: -.19, .04), but remained significant for females (-.13, CI: -.24, -.02; Figure 1).

Individuals with more bonds to flock mates (higher degree) had a higher number of fledglings (F: .19, CI: .07, .31; M: .18, CI: .06, .29), but this effect was not significant after controlling for spatial autocorrelation (F: .10, CI: -.03, .23; M: .09, CI: -.04, .22). Individuals with more bonds to flock mates had later lay dates, and this effect remained significant after controlling for breeding spatial autocorrelation (F: .10, CI: .03, .18; M: .10, CI: .03, .18; Figure 1).

Individuals with more winter spatial associates had smaller clutches, but this effect was only significant after controlling for breeding spatial autocorrelation (F: -.14, CI: -.26, -.03; M: - .14, CI: -.26, -.03; Figure 1). Although it was retained in the models, there were no significant effects of density on lay date (F: .06, CI: -.01, .13; M: .05, CI: -.02, .13).

### Non-social spatial drivers

Individuals who bred in areas with a better habitat quality had earlier lay dates (F: -.09, CI: - .16, -.02; M: -.09, CI: -.16, -.02) and larger clutches (F: .11, CI: .01, .21; M: 11, CI: .01, .21; Figure 3), but this effect was no longer significant after controlling for breeding spatial autocorrelation.

The spatial distribution of each of the response (fitness) variables is graphically illustrated by projecting the SPDE random effect onto a two-dimensional plane (Figure 2). Overall, the southern part of the woods has lower reproductive success than the northern part, and lay dates are the latest in the central portion of the woods. Adding the SPDE random effect (accounting for breeding spatial autocorrelation) improves all of the models substantially, with the exception of the binary success model (Table 1).

**Figure 2.**
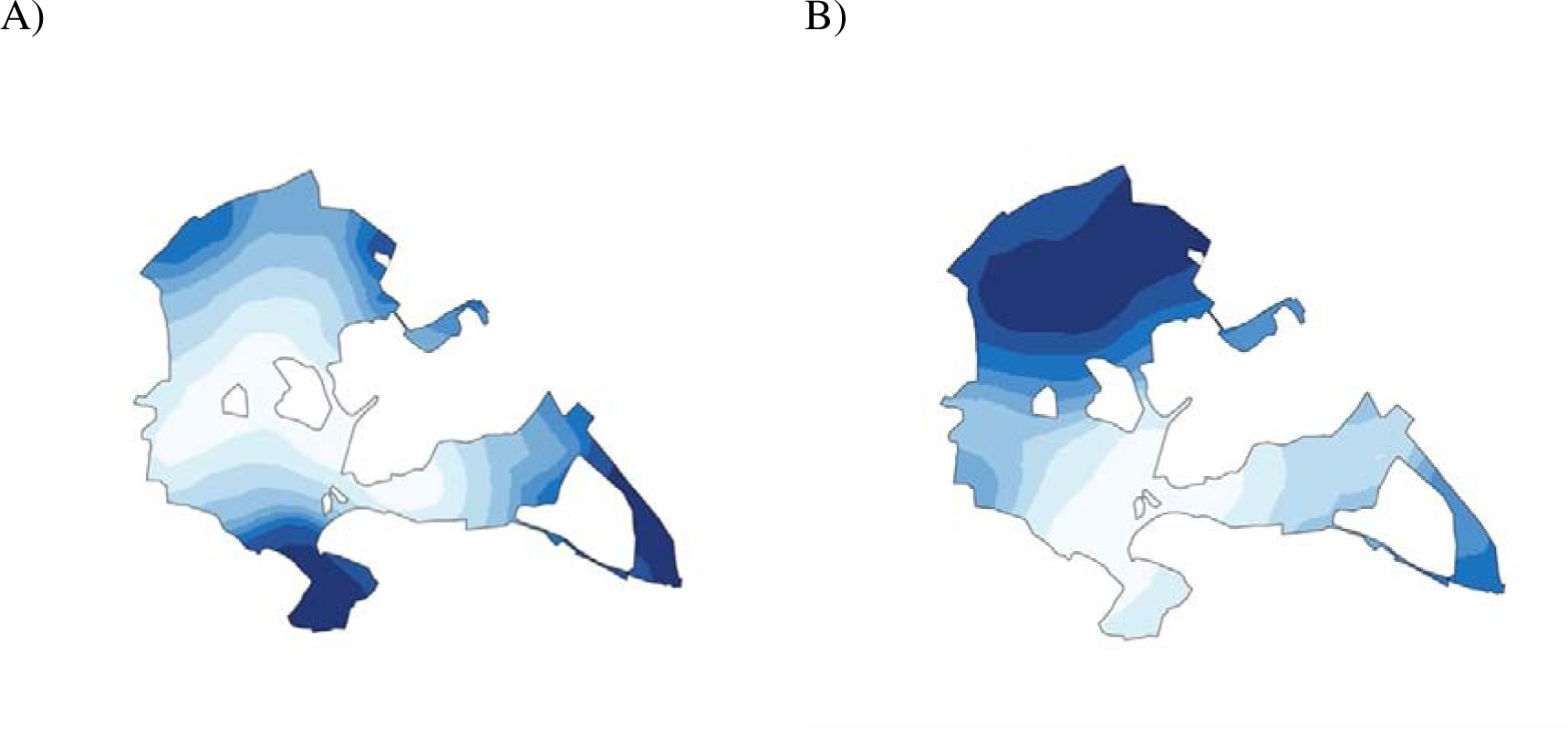
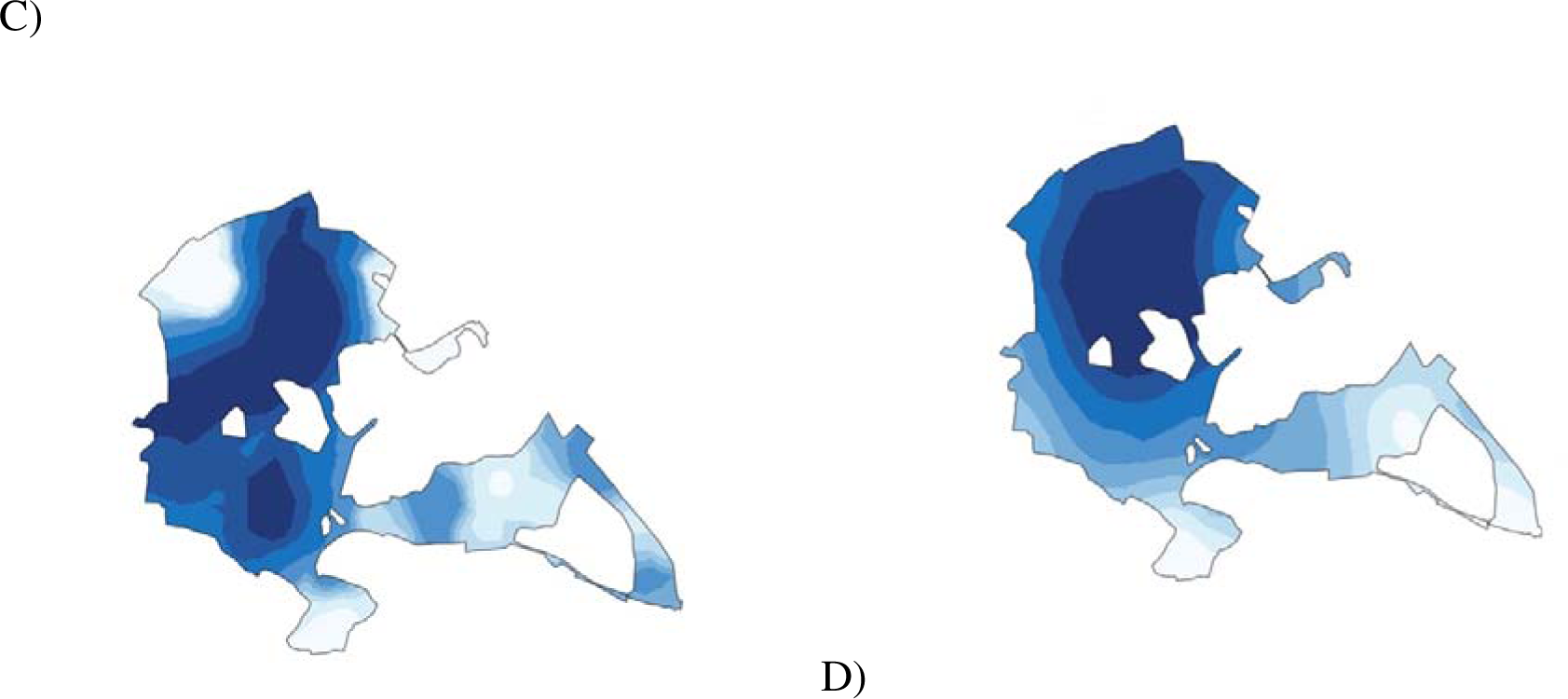
- 2D projection of the spatial distribution of the z-scores for each fitness variable (for females), when accounting for both the fixed and random effects in each model. For a) lay date, b) clutch size, c) mean chick weight, and d) number of fledglings. Darker shading denotes higher success.

**Table 1.** - DIC for base and SPDE models

| Model | Base | SPDE | DeltaDIC |
| --- | --- | --- | --- |
| Lay date | 570.145 | 550.329 | -19.816 |
| Binary success | 1046.171 | 1045.140 | -1.030 |
| Clutch size | 933.781 | 906.790 | -26.991 |
| Mean chick weight | 901.932 | 880.485 | -21.447 |
| Number of fledglings | 1028.322 | 1007.455 | -20.867 |

### Other non-social drivers

Year had an influence on all of the variables except for mean chick weight (see Figure 3). There were no significant differences in terms of numeric age. However, juveniles had significantly later lay dates than adults (F: .35, CI: .21, .48; M: .35, CI: .17, .54). All social and non-social drivers were included in the same models (Table 2).

**Figure 3.**
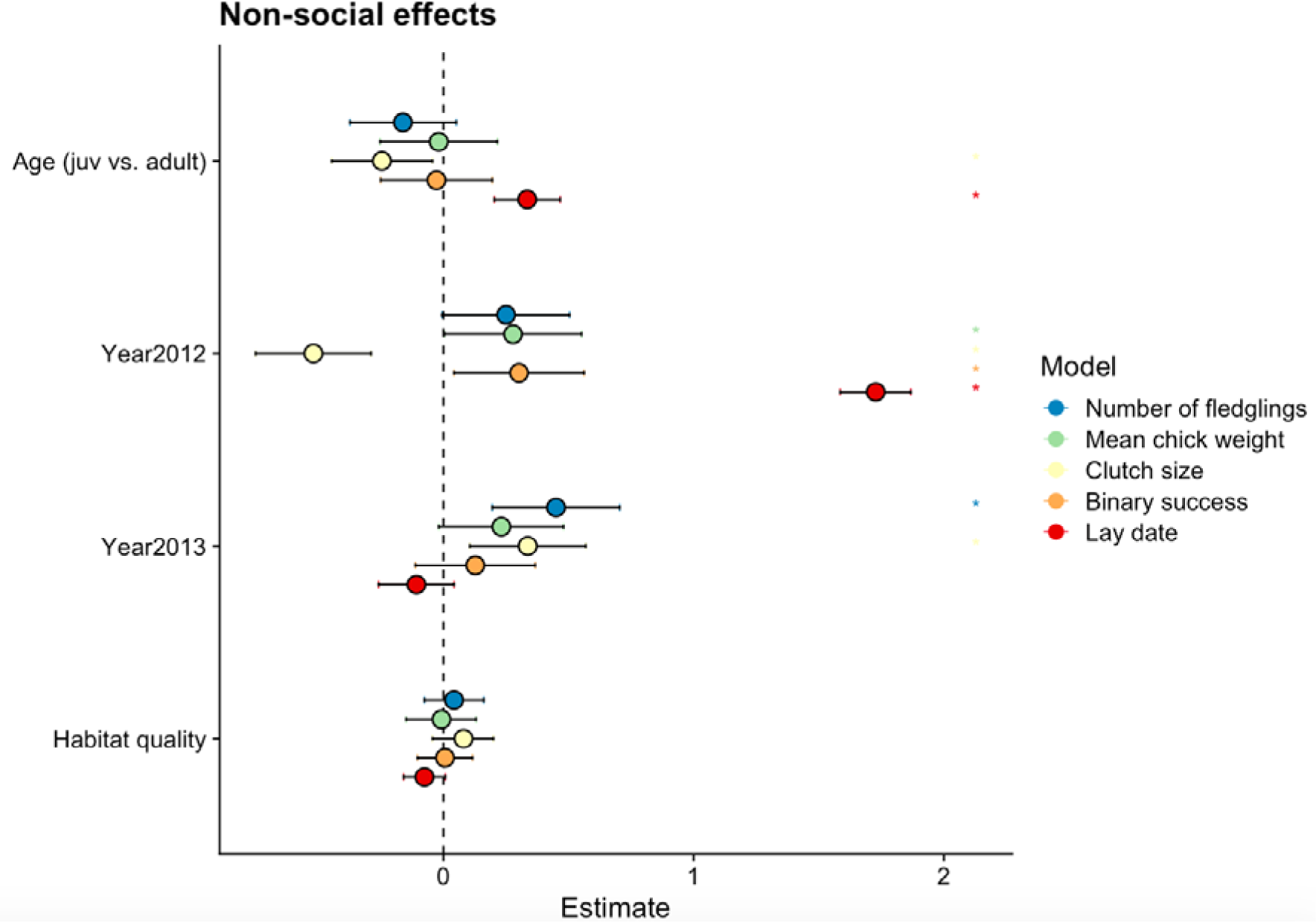
- Summary of the non-social effects with the SPDE effect to control for spatial autocorrelation. Points represent the estimate for each effect that was retained in the model selection process and error bars denote 95% credibility intervals

**Table 2.**
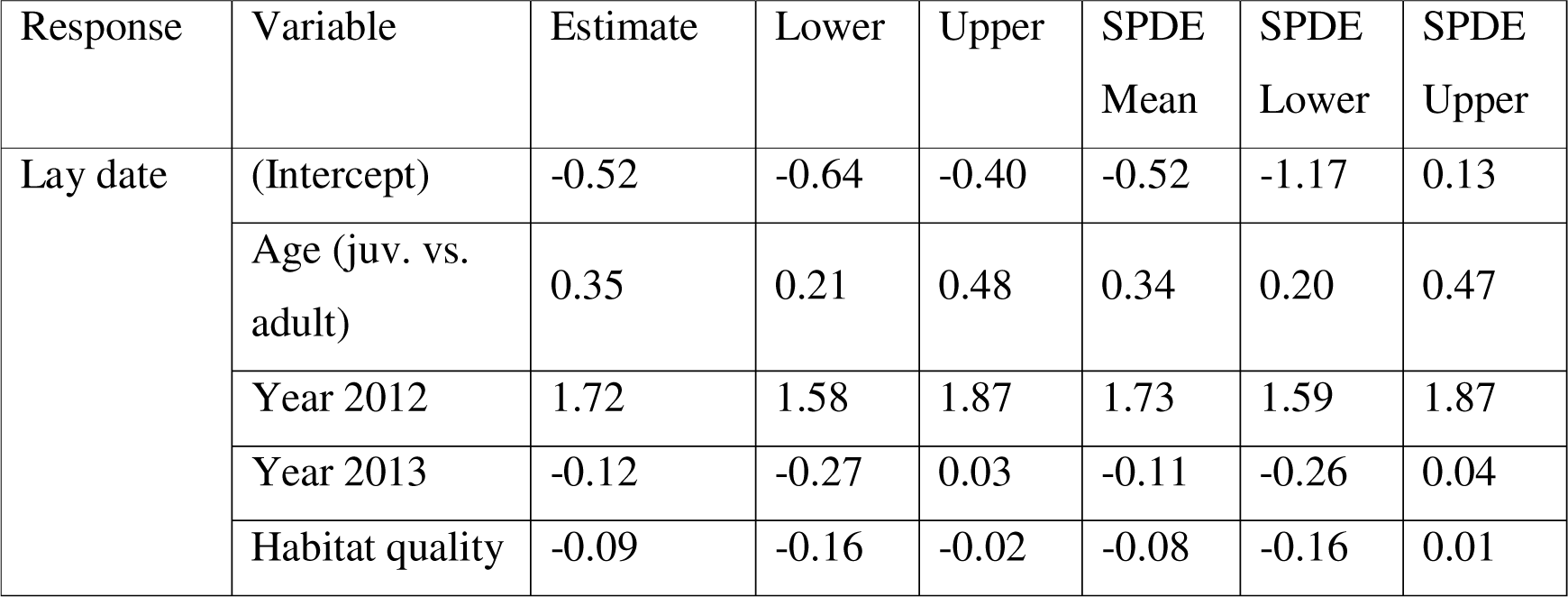
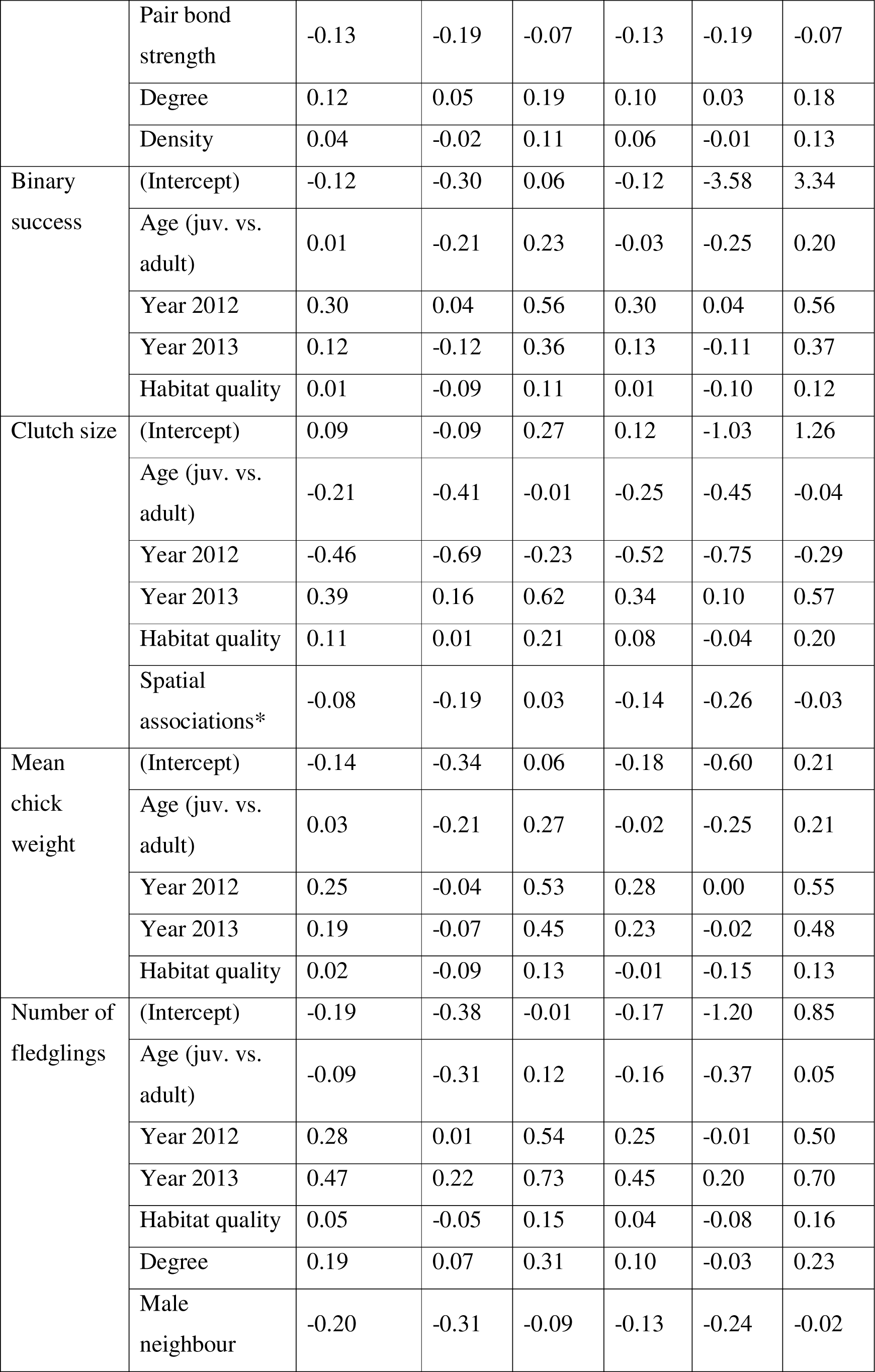
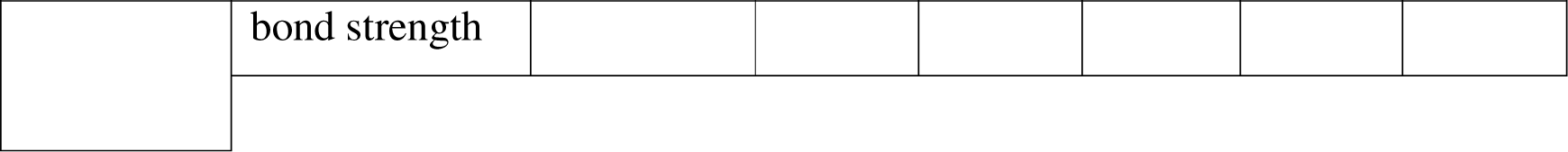
- model statistics for female main models (same information for males can be found in Supplementary Table 1); bold variables are significant in both models, italic variables are only significant before controlling for breeding spatial autocorrelation, * variables are only significant after controlling for breeding spatial autocorrelation

## Discussion

Although individuals exist in complex multi-level societies, much work exploring the relationship between sociality and fitness has focused on a single type of dyadic bond. Using three years of behavioural and reproductive data in wild songbirds, we demonstrate the potential fitness consequences of four types of dyadic relationships that are found within multilevel societies. Individuals with a stronger bond to their pair mate had earlier lay dates, consistent with previous findings (Culina *et al*. 2020). Although more social individuals (higher winter network degree) had later lay dates, they also had a higher number of fledglings, suggesting that social associations with flock mates influence fitness outcomes in multiple dimensions, rather than simply improving fitness overall. When controlling for other social and spatial effects, bonds with breeding neighbours were not additionally beneficial. In fact, having stronger bonds with neighbouring males was associated with having slightly fewer fledglings. Finally, individuals with more winter spatial associates had smaller clutch sizes – suggesting that it is important to control for spatial relationships when considering other, more traditionally studied, aspects of sociality. Accounting for breeding position spatial autocorrelation in fitness outcomes improved nearly all of the models, highlighting the importance of accounting for spatial effects.

Our results demonstrate support for a consistent finding in this population - the importance of the pair bond in determining lay dates, with individuals that met earlier in the winter (Culina *et al*. 2020) or had bred together in the past (Gokcekus *et al*. 2023) having earlier lay dates. Here, we show this relationship still holds even when simultaneously considering other kinds of social links. Historically, many external factors have been used to predict lay dates (Perrins & McCleery 1989), including temperature (Noordwijk *et al*. 1995; Visser *et al*. 1998), habitat quality (Wilkin *et al*. 2007), caterpillar growth (Visser *et al*. 2006), and presence of parasites (Oppliger *et al*. 1994) among others. Yet the strength of the pair bond seems to consistently be an important predictor that is not usually examined. Lay date has often been considered as an individual trait, but these results suggest that it may also be a social trait (and therefore dependent on others in the social system; see discussion below on flock mate relations and lay date)(Evans *et al*. 2020).

Previous studies in this population have shown that individuals breed close to their highly associated former flock mates (Firth & Sheldon 2016) and that neighbours who are socially familiar (due to year-to-year territory sharing) have higher fitness (Gokcekus *et al*. 2023). However, when controlling for other types of dyadic bonds, having strong winter bonds to subsequent breeding neighbours was not actually beneficial in this sample. Females with stronger bonds to neighbouring males had fewer fledglings. It is possible that tolerating males may actually come at a cost if it leads to having fewer resources. Additionally, if females with stronger bonds to male neighbours are more likely to have extra-pair copulations, males may subsequently provide less parental care in the nest or spend more time investing in other females (Eliassen & Jørgensen 2014; Magrath & Komdeur 2003).

This finding on breeding neighbours raises the possibility that the previously observed positive effect of neighbour familiarity (Gokcekus *et al*. 2023; Grabowska-Zhang *et al*. 2012c) is actually due to general sociality and/or early establishing of territories and not the result of cooperation that arises due to specific bonds with particular individuals. However, it is also possible that breeding neighbour familiarity is most important for an aspect of fitness that we cannot measure: whether individuals are able to establish a territory at all. In support of this possibility, a previous study on this population showed that immigrants are more likely to fail in establishing a territory (and therefore establishing a breeding attempt) than integrated members (Kidd *et al*. 2015). Furthermore, it is possible that individuals that are familiar over multiple years are easier to identify because they become associated with territory boundaries. If there is a limit on the number of individuals that great tits can identify, it is possible that having stronger bonds with neighbours is not as beneficial as having neighbours that are familiar over the years simply because birds are more likely to be aware of this between-year familiarity (Gokcekus *et al*. 2021).

Individuals with a higher degree during the winter have more fledglings, which is in line with the general idea that being more social comes with fitness benefits (Snyder-Mackler *et al*. 2020). However, this effect is weakened when considering breeding location spatial effects, raising the possibility that part of this relationship may be explained by higher-quality areas supporting stronger bonds. On the other hand, individuals with higher degree also had later lay dates, which may represent that dedicating social time to other flockmates trades off against the social bond with their pair mate and thus potentially pushes back the lay date. When considering winter spatial relations, we find that individuals with more winter spatial associates have smaller clutches. Individuals who spent the winter in higher-density areas had slightly later lay dates but this effect was not significant. As resources are provided only on weekends at these feeding stations, it is unlikely that the reduced clutch sizes are the result of resource competition at the feeders themselves; instead, females who spend the winter in areas with more conspecifics might anticipate greater competition and lay smaller clutches accordingly to avoid overstretching the available resources. This result also highlights the importance of explicitly testing and controlling for spatial associations when investigating more apparent measures of sociality.

In this multi-level society, four types of social relationships measured across seasons influenced breeding success when tracked over multiple years. However, future experimental work is likely necessary to understand the causation behind these effects. For example, sociality’s influence on reproductive success could be investigated by manipulating the ability of some individuals to form social bonds. Detailed data on different kinds of social associations across multiple circumstances and years is necessary for a more thorough knowledge of the fitness consequences of sociality within multi-level societies. Disentangling the effects of different types of dyadic bonds within such societies and separating them from spatial patterns is important for gaining a more thorough understanding of the factors that drive the evolution of social stability within populations.

## Acknowledgements

This work was supported by grants from BBSRC (BB/L006081/1, BB/S009752/1), ERC (AdG250164), and NERC (NE/K006274/1, NE/S010335/1, NE/V013483/1) and the Wild Animal Initiative (C-2023-00057).

## Supplementary material

**Supplementary Figure 1.**
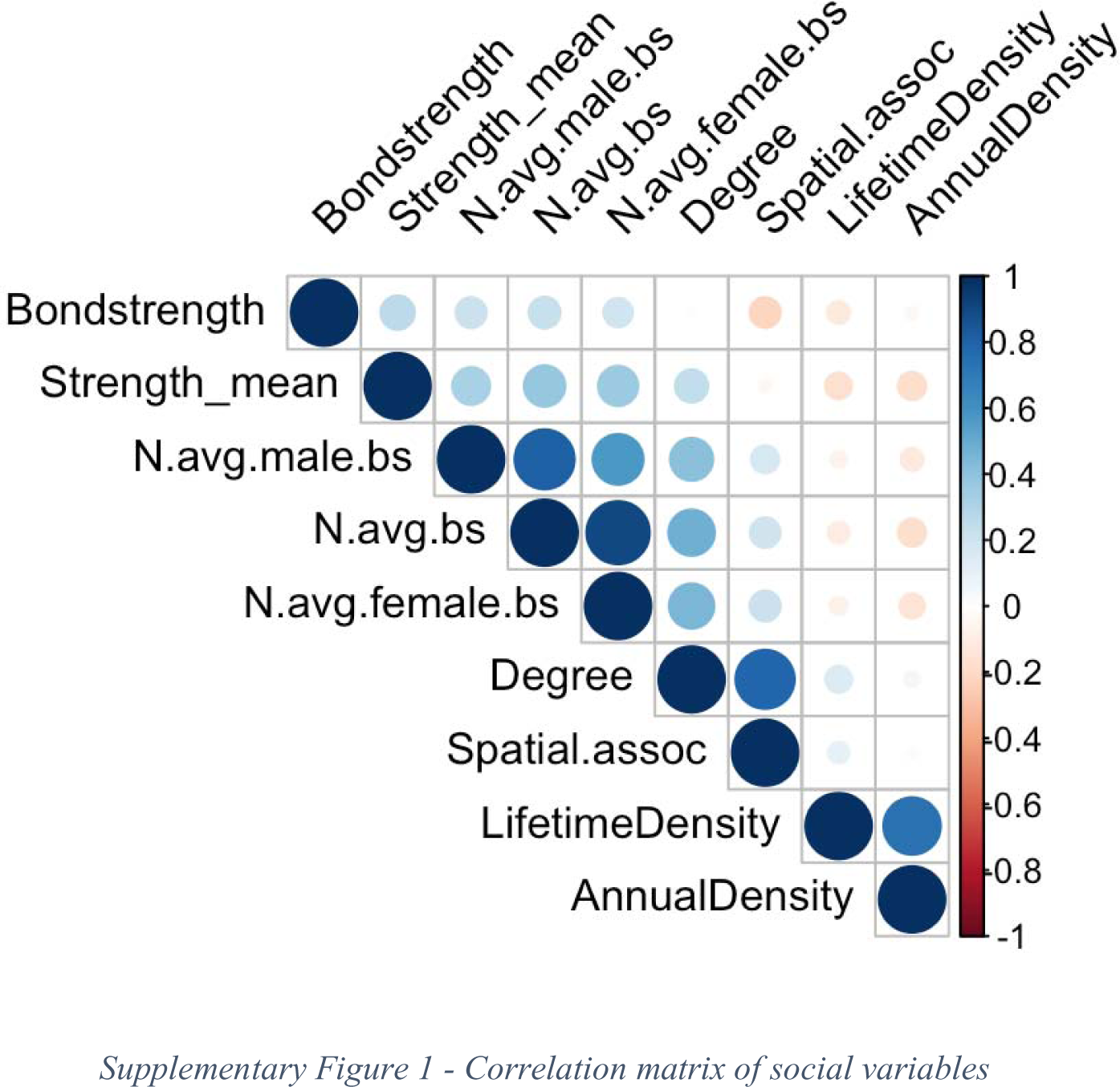
- Correlation matrix of social variables

**Supplementary Figure 2.**
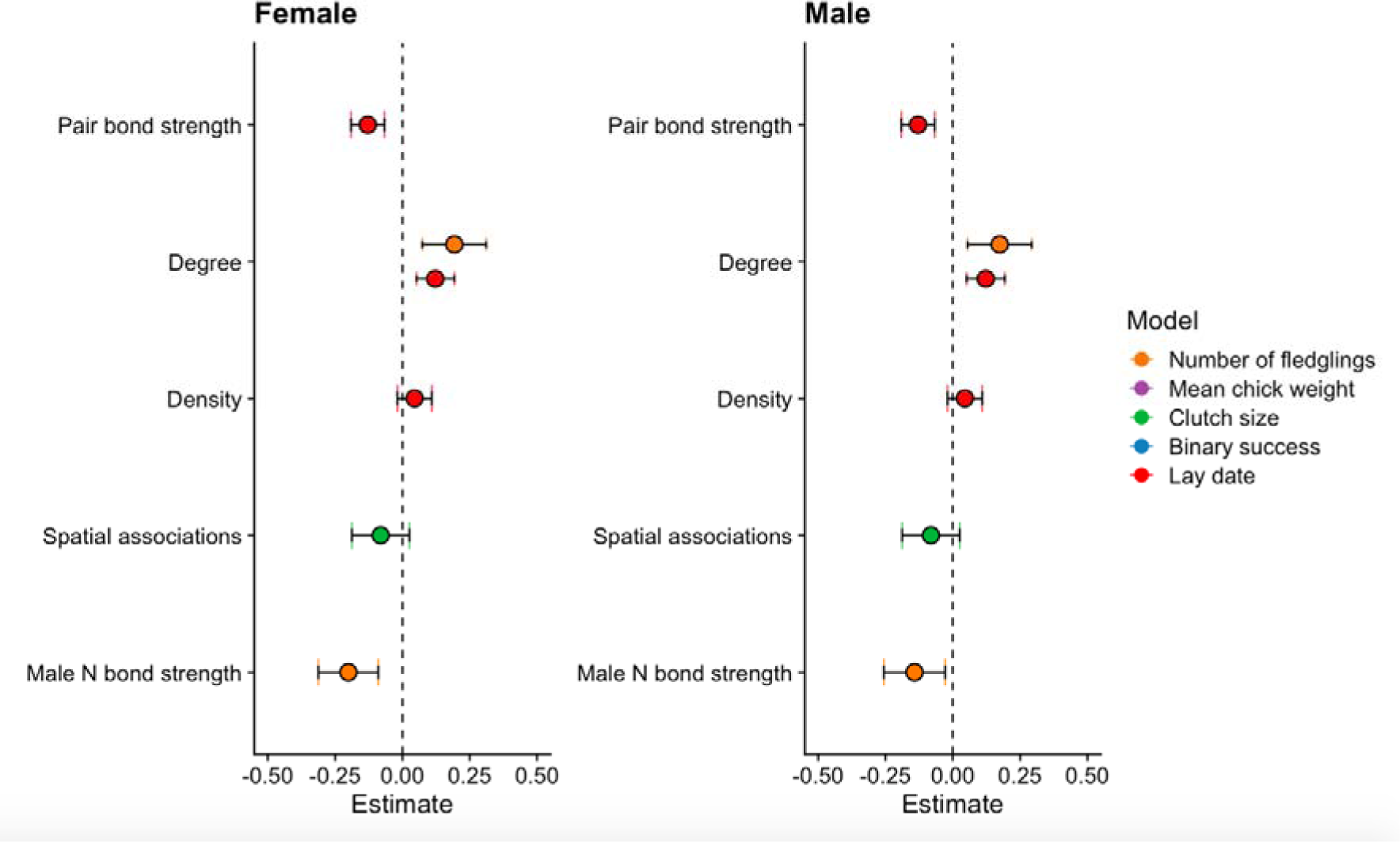
- model results (only social effects) without SPDE spatial autocorrelation correction

**Supplementary Figure 3.**
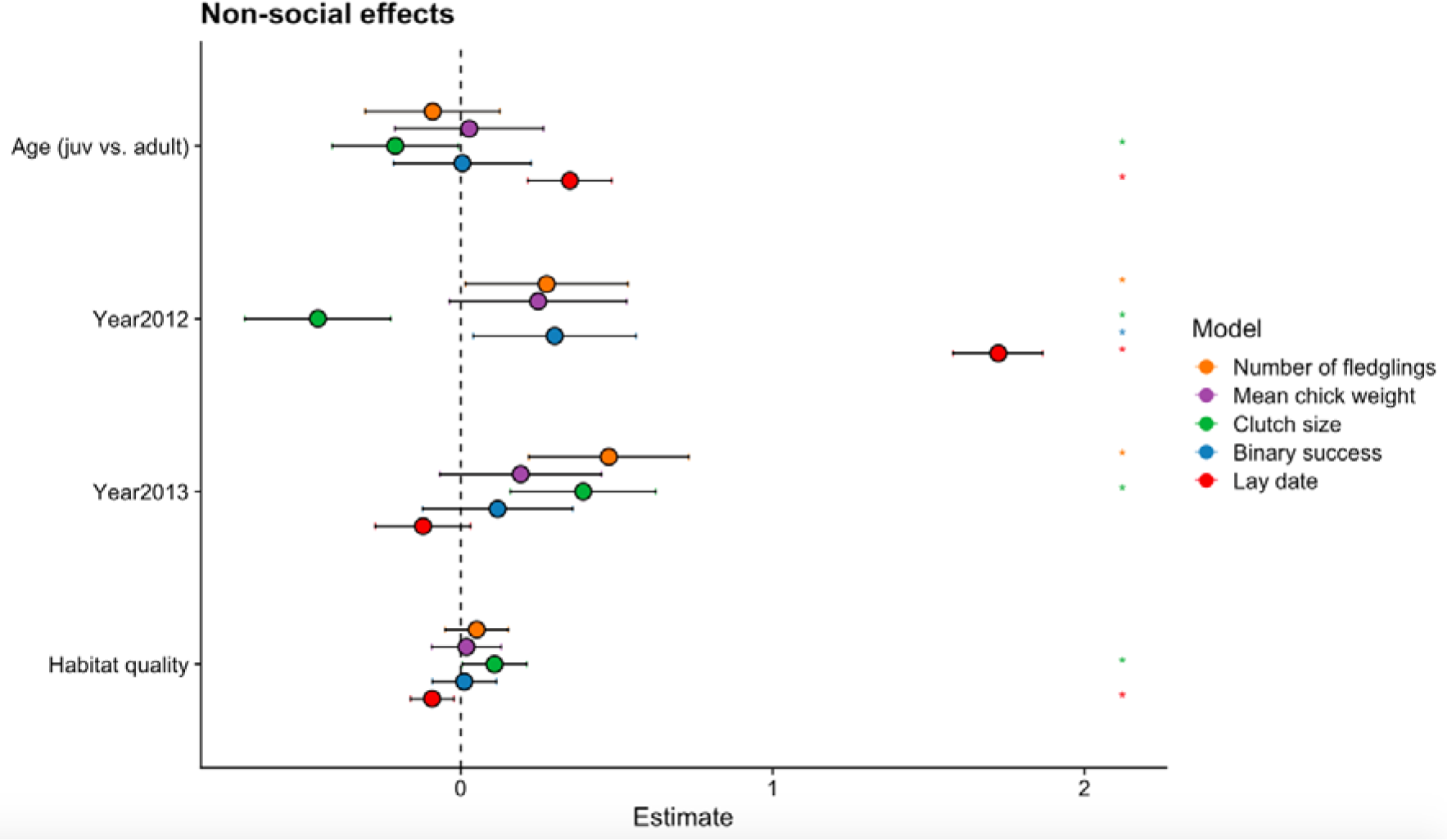
- model results (only non-social effects without SPDE spatial autocorrelation correction

**Supplementary Table 1.** - Male model results

| Response | Variable | Estimate | Lower | Upper | SPDE<br>Mean | SPDE<br>Lower | SPDE<br>Upper |
| --- | --- | --- | --- | --- | --- | --- | --- |
| Lay date | (Intercept) | -0.52 | -0.64 | -0.40 | -0.52 | -1.24 | 0.18 |
|  | Age (juv. vs. adult) | 0.35 | 0.17 | 0.54 | 0.37 | 0.19 | 0.55 |
|  | Year 2012 | 1.72 | 1.58 | 1.87 | 1.73 | 1.58 | 1.86 |
|  | Year 2013 | -0.12 | -0.28 | 0.04 | -0.12 | -0.28 | 0.04 |
|  | Habitat quality | -0.09 | -0.16 | -0.02 | -0.08 | -0.16 | 0.01 |
|  | Pair bond strength | -0.13 | -0.19 | -0.07 | -0.13 | -0.19 | -0.07 |
|  | Degree | 0.12 | 0.05 | 0.19 | 0.10 | 0.03 | 0.18 |
|  | Density | 0.04 | -0.02 | 0.11 | 0.05 | -0.02 | 0.13 |
| Binary success | (Intercept) | -0.11 | -0.30 | 0.09 | -0.11 | -3.21 | 2.99 |
|  | Age (juv. vs. adult) | -0.04 | -0.36 | 0.28 | -0.07 | -0.39 | 0.24 |
|  | Year 2012 | 0.30 | 0.04 | 0.57 | 0.30 | 0.04 | 0.56 |
|  | Year 2013 | 0.13 | -0.12 | 0.37 | 0.14 | -0.11 | 0.38 |
|  | Habitat quality | 0.01 | -0.09 | 0.12 | 0.01 | -0.10 | 0.12 |
| Clutch size | (Intercept) | 0.08 | -0.11 | 0.26 | 0.11 | -0.87 | 1.09 |
|  | Age (juv. vs. adult) | -0.17 | -0.45 | 0.11 | -0.21 | -0.49 | 0.07 |
|  | Year 2012 | -0.46 | -0.70 | -0.23 | -0.52 | -0.76 | -0.29 |
|  | Year 2013 | 0.38 | 0.14 | 0.62 | 0.33 | 0.09 | 0.56 |
|  | Habitat quality | 0.11 | 0.01 | 0.21 | 0.08 | -0.05 | 0.20 |
|  | Spatial associations* | -0.08 | -0.19 | 0.03 | -0.14 | -0.26 | -0.03 |
| Mean chick weight | (Intercept) | -0.18 | -0.40 | 0.03 | -0.23 | -0.66 | 0.17 |
|  | Age (juv. vs. adult) | 0.17 | -0.18 | 0.52 | 0.17 | -0.17 | 0.51 |
|  | Year 2012 | 0.24 | -0.04 | 0.53 | 0.27 | -0.00 | 0.54 |
|  | Year 2013 | 0.17 | -0.10 | 0.43 | 0.20 | -0.06 | 0.45 |
|  | Habitat quality | 0.02 | -0.09 | 0.13 | -0.01 | -0.15 | 0.13 |
| Number of fledglings | (Intercept) | -0.17 | -0.38 | 0.03 | -0.18 | -1.24 | 0.88 |
|  | Age (juv. vs. adult) | -0.09 | -0.41 | 0.24 | -0.11 | -0.42 | 0.21 |
|  | Year 2012 | 0.24 | -0.02 | 0.51 | 0.22 | -0.04 | 0.47 |
|  | Year 2013 | 0.45 | 0.20 | 0.71 | 0.43 | 0.18 | 0.68 |
|  | Habitat quality | 0.06 | -0.04 | 0.16 | 0.06 | -0.06 | 0.18 |
|  | Male neighbour bond strength | 0.18 | 0.06 | 0.29 | 0.09 | -0.04 | 0.22 |
|  | Degree | -0.14 | -0.26 | -0.03 | -0.07 | -0.19 | 0.04 |

## References

1. Adams, E.S. (2001). Approaches to the study of territory size and shape. Annu Rev Ecol Syst.

2. Alberts, S.C. (2019). Social influences on survival and reproduction: Insights from a long-term study of wild baboons. Journal of Animal Ecology, 88, 47–66.

3. Albery, G.F., Becker, D.J., Kenyon, F., Nussey, D.H. & Pemberton, J.M. (2019). The fine-scale landscape of immunity and parasitism in a wild ungulate population. Integr Comp Biol, 59, 1165–1175.

4. Albery, G.F., Clutton-Brock, T.H., Morris, A., Morris, S., Pemberton, J.M., Nussey, D.H., et al. (2022). Ageing red deer alter their spatial behaviour and become less social. Nat Ecol Evol, 6, 1231–1238.

5. Albery, G.F., Morris, A., Morris, S., Pemberton, J.M., CluttonlBrock, T.H., Nussey, D.H., et al. (2021). Multiple spatial behaviours govern social network positions in a wild ungulate. Ecol Lett, 24, 676–686.

6. Albery, G.F., Newman, C., Bright Ross, J., Macdonald, D.W., Bansal, S. & Buesching, C.D. (2020). Negative density-dependent parasitism in a group-living carnivore. Proceedings of the Royal Society B: Biological Sciences, 287, 20202655.

7. Aplin, L.M., Farine, D.R., Morand-Ferron, J., Cole, E.F., Cockburn, A. & Sheldon, B.C. (2013). Individual personalities predict social behaviour in wild networks of great tits (Parus major). Ecol Lett, 16, 1365–1372.

8. Ausband, D.E. (2019). Pair bonds, reproductive success, and rise of alternate mating strategies in a social carnivore. Behavioral Ecology, 30.

9. Bebbington, K., Kingma, S.A., Fairfield, E.A., Dugdale, H.L., Komdeur, J., Spurgin, L.G., et al. (2017). Kinship and familiarity mitigate costs of social conflict between Seychelles warbler neighbors. Proc Natl Acad Sci U S A, 114, E9036–E9045.

10. Beck, K.B., Farine, D.R. & Kempenaers, B. (2020). Winter associations predict social and extra-pair mating patterns in a wild songbird. Proc Biol Sci, 287, 20192606.

11. Beletsky, L.D. & Orians, G.H. (1989). Familiar neighbors enhance breeding success in birds. *Proceedings - National Academy of Sciences*, USA, 86, 7933–7936.

12. Berghänel, A., Ostner, J., Schröder, U. & Schülke, O. (2011). Social bonds predict future cooperation in male Barbary macaques, Macaca sylvanus. Anim Behav, 81, 1109–1116.

13. Bond, M.L., Lee, D.E., Farine, D.R., Ozgul, A. & König, B. (2021). Sociability increases survival of adult female giraffes. Proceedings of the Royal Society B: Biological Sciences, 288, 20202770.

14. Booksmythe, I., Hayes, C., Jennions, M.D. & Backwell, P.R.Y. (2012). The effects of neighbor familiarity and size on cooperative defense of fiddler crab territories. Behavioral Ecology, 23.

15. Bray, J., Feldblum, J.T. & Gilby, I.C. (2021). Social bonds predict dominance trajectories in adult male chimpanzees. Anim Behav.

16. Cameron, E.Z., Setsaas, T.H. & Linklater, W.L. (2009). Social bonds between unrelated females increase reproductive success in feral horses. Proc Natl Acad Sci U S A, 106, 13850–13853.

17. Campbell, L.A.D., Tkaczynski, P.J., Lehmann, J., Mouna, M. & Majolo, B. (2018). Social thermoregulation as a potential mechanism linking sociality and fitness: Barbary macaques with more social partners form larger huddles. Sci Rep, 8.

18. Cheney, D.L., Silk, J.B. & Seyfarth, R.M. (2016). Network connections, dyadic bonds and fitness in wild female baboons. R Soc Open Sci, 3.

19. Clutton-Brock, T.H. & Sheldon, B.C. (2010). Individuals and populations: The role of long-term, individual-based studies of animals in ecology and evolutionary biology. Trends Ecol Evol, 25, 562–573.

20. Culina, A., Firth, J.A. & Hinde, C.A. (2020). Familiarity breeds success: Pairs that meet earlier experience increased breeding performance in a wild bird population: Pairs that meet earlier do better. Proceedings of the Royal Society B: Biological Sciences, 287, 20201554.

21. Culina, A., Lachish, S. & Sheldon, B.C. (2015a). Evidence of a link between survival and pair fidelity across multiple tit populations. J Avian Biol, 46, 507–515.

22. Culina, A., Radersma, R. & Sheldon, B.C. (2015b). Trading up: The fitness consequences of divorce in monogamous birds. Biological Reviews, 90, 1015–1034.

23. Eliassen, S. & Jørgensen, C. (2014). Extra-pair mating and evolution of cooperative neighbourhoods. PLoS One, 9.

24. Ellis, S., Snyder-Mackler, N., Ruiz-Lambides, A., Platt, M.L. & Brent, L.J.N. (2019). Deconstructing sociality: The types of social connections that predict longevity in a group-living primate. Proceedings of the Royal Society B: Biological Sciences, 286.

25. Evans, S.R., Postma, E. & Sheldon, B.C. (2020). It takes two: Heritable male effects on reproductive timing but not clutch size in a wild bird population*. Evolution (N Y), 74.

26. Farine, D.R., Firth, J.A., Aplin, L.M., Crates, R.A., Culina, A., Garroway, C.J., et al. (2015). The role of social and ecological processes in structuring animal populations: A case study from automated tracking of wild birds. R Soc Open Sci, 2.

27. Farine, D.R. & Sheldon, B.C. (2015). Selection for territory acquisition is modulated by social network structure in a wild songbird. J Evol Biol, 28, 547–556.

28. Fayet, A.L., Shoji, A., Freeman, R., Perrins, C.M. & Guilford, T. (2017). Within-pair similarity in migration route and female winter foraging effort predict pair breeding performance in a monogamous seabird. Mar Ecol Prog Ser, 569.

29. Feldblum, J.T., Krupenye, C., Bray, J., Pusey, A.E. & Gilby, I.C. (2021). Social bonds provide multiple pathways to reproductive success in wild male chimpanzees. iScience, 24, 102864.

30. Firth, J.A., Cole, E.F., Ioannou, C.C., Quinn, J.L., Aplin, L.M., Culina, A., et al. (2018a). Personality shapes pair bonding in a wild bird social system. Nat Ecol Evol, 2, 1696– 1699.

31. Firth, J.A. & Sheldon, B.C. (2016). Social carry-over effects underpin trans-seasonally linked structure in a wild bird population. Ecol Lett, 19, 1324–1332.

32. Firth, J.A., Sheldon, B.C. & Farine, D.R. (2016). Pathways of information transmission among wild songbirds follow experimentally imposed changes in social foraging structure. Biol Lett, 12.

33. Firth, J.A., Verhelst, B.L., Crates, R.A., Garroway, C.J. & Sheldon, B.C. (2018b). Spatial, temporal and individual-based differences in nest-site visits and subsequent reproductive success in wild great tits. J Avian Biol, 49.

34. Firth, J.A., Voelkl, B., Crates, R.A., Aplin, L.M., Biro, D., Croft, D.P., et al. (2017). Wild birds respond to flockmate loss by increasing their social network associations to others. Proceedings of the Royal Society B: Biological Sciences, 284.

35. Firth, J.A., Voelkl, B., Farine, D.R. & Sheldon, B.C. (2015). Experimental evidence that social relationships determine individual foraging behavior. Current Biology, 25, 3138– 3143.

36. Fisher, D.N., Kilgour, R.J., Eiracusa, E.R., Foote, J.R., Hobson, E.A., Montiglio, P.-O., et al. (2021). Anticipated effects of abiotic environmental change on intraspecific social interactions. Biological Reviews, 44.

37. Gokcekus, S., Firth, J.A., Regan, C., Cole, E.F., Sheldon, B.C. & Albery, G.F. (2023). Social familiarity and spatially variable environments independently determine reproductive fitness in a wild bird. American Naturalist.

38. Gokcekus, S., Firth, J.A., Regan, C. & Sheldon, B.C. (2021). Recognising the key role of individual recognition in social networks. Trends Ecol Evol, 36, 1024–1035.

39. Grabowska-Zhang, A.M., Sheldon, B.C. & Hinde, C.A. (2012a). Long-term familiarity promotes joining in neighbour nest defence. Biol Lett, 8, 544–546.

40. Grabowska-Zhang, A.M., Wilkin, T.A. & Sheldon, B.C. (2012b). Effects of neighbor familiarity on reproductive success in the great tit (Parus major). Behavioral Ecology, 23, 322–333.

41. Grabowska-Zhang, A.M., Wilkin, T.A. & Sheldon, B.C. (2012c). Effects of neighbor familiarity on reproductive success in the great tit (Parus major). Behavioral Ecology, 23, 322–333.

42. Griffith, S.C., Pryke, S.R. & Buttemer, W.A. (2011). Constrained mate choice in social monogamy and the stress of having an unattractive partner. Proceedings of the Royal Society B: Biological Sciences, 278.

43. Grueter, C.C., Qi, X., Li, B. & Li, M. (2017). Multilevel societies. Current Biology, 27, R984–R986.

44. Grueter, C.C., Qi, X., Zinner, D., Bergman, T., Li, M., Xiang, Z., et al. (2020). Multilevel Organisation of Animal Sociality. Trends Ecol Evol.

45. He, P., Maldonado-Chaparro, A.A. & Farine, D.R. (2019). The role of habitat configuration in shaping social structure: a gap in studies of animal social complexity. Behav Ecol Sociobiol, 73.

46. Ihle, M., Kempenaers, B. & Forstmeier, W. (2015). Fitness Benefits of Mate Choice for Compatibility in a Socially Monogamous Species. PLoS Biol, 13.

47. Kidd, L.R., Sheldon, B.C., Simmonds, E.G. & Cole, E.F. (2015). Who escapes detection? Quantifying the causes and consequences of sampling biases in a long-term field study. Journal of Animal Ecology, 84, 1520–1529.

48. Kohn, G.M. (2017). Friends give benefits: autumn social familiarity preferences predict reproductive output. Anim Behav, 132, 201–208.

49. Liebgold, E.B. & Cabe, P.R. (2008). Familiarity with adults, but not relatedness, affects the growth of juvenile red-backed salamanders (Plethodon cinereus). Behav Ecol Sociobiol, 63, 277–284.

50. Magrath, M.J.L. & Komdeur, J. (2003). Is male care compromised by additional mating opportunity? Trends Ecol Evol, 18, 424–430.

51. Massen, J.J.M., Sterck, E.H.M. & De Vos, H. (2010). Close social associations in animals and humans: Functions and mechanisms of friendship. Behaviour, 147, 1379–1412.

52. McFarland, R., Murphy, D., Lusseau, D., Henzi, S.P., Parker, J.L., Pollet, T. V., et al. (2017). The ‘strength of weak ties’ among female baboons: fitness-related benefits of social bonds. Anim Behav, 126, 101–106.

53. Menz, C.S., Carter, A.J., Best, E.C., Freeman, N.J., Dwyer, R.G., Blomberg, S.P., et al. (2020). Higher sociability leads to lower reproductive success in female kangaroos: Sociability and fitness in kangaroos. R Soc Open Sci, 7.

54. Micheletta, J., Waller, B.M., Panggur, M.R., Neumann, C., Duboscq, J., Agil, M., et al. (2012). Social bonds affect anti-predator behaviour in a tolerant species of macaque, Macaca nigra. Proceedings of the Royal Society B: Biological Sciences, 279.

55. Müller, C.A. & Manser, M.B. (2007). “Nasty neighbours” rather than “dear enemies” in a social carnivore. Proceedings of the Royal Society B: Biological Sciences, 274.

56. Noordwijk, A.J. Van, McCleery, R.H. & Perrins, C.M. (1995). Selection for the Timing of Great Tit Breeding in Relation to Caterpillar Growth and Temperature. J Anim Ecol, 64.

57. Oppliger, A., Richner, H. & Christe, P. (1994). Effect of an ectoparasite on lay date, nest-site choice, desertion, and hatching success in the great tit (Pants major). Behavioral Ecology, 5.

58. Papageorgiou, D., Christensen, C., Gall, G.E.C., Klarevas-Irby, J.A., Nyaguthii, B., Couzin, I.D., et al. (2019). The multilevel society of a small-brained bird. Current Biology.

59. Papageorgiou, D. & Farine, D.R. (2021). Multilevel Societies in Birds. Trends Ecol Evol.

60. Perrins, C.M. (1965). Population Fluctuations and Clutch-Size in the Great Tit, Parus major L. J Anim Ecol, 34.

61. Perrins, C.M. & McCleery, R.H. (1989). Laying date and clutch size in the great tit. Wilson Bulletin, 101.

62. Van De Pol, M., Heg, D., Bruinzeel, L.W., Kuijper, B. & Verhulst, S. (2006). Experimental evidence for a causal effect of pair-bond duration on reproductive performance in oystercatchers (Haematopus ostralegus). Behavioral Ecology, 17.

63. Psorakis, I., Roberts, S.J., Rezek, I. & Sheldon, B.C. (2012). Inferring social network structure in ecological systems from spatiotemporal data streams. J R Soc Interface.

64. Riehl, C. & Strong, M.J. (2018). Stable social relationships between unrelated females increase individual fitness in a cooperative bird. Proceedings of the Royal Society B: Biological Sciences, 285.

65. Rue, H., Martino, S. & Chopin, N. (2009). Approximate Bayesian inference for latent Gaussian models by using integrated nested Laplace approximations. J R Stat Soc Series B Stat Methodol, 71, 319–392.

66. Sabol, A.C., Lambert, C.T., Keane, B., Solomon, N.G. & Dantzer, B. (2020). How does individual variation in sociality influence fitness in prairie voles? Anim Behav, 163.

67. Samuni, L., Crockford, C. & Wittig, R.M. (2021). Group-level cooperation in chimpanzees is shaped by strong social ties. Nat Commun, 12.

68. Samuni, L., Preis, A., Mielke, A., Deschner, T., Wittig, R.M. & Crockford, C. (2018). Social bonds facilitate cooperative resource sharing in wild chimpanzees. Proceedings of the Royal Society B: Biological Sciences, 285, 20181643.

69. Sánchez-Macouzet, O., Rodríguez, C. & Drummond, H. (2014). Better stay together: Pair bond duration increases individual fitness independent of age-related variation. Proceedings of the Royal Society B: Biological Sciences, 281.

70. Schlicht, L., Valcu, M. & Kempenaers, B. (2014). Thiessen polygons as a model for animal territory estimation. Ibis, 156, 215–219.

71. Schülke, O., Bhagavatula, J., Vigilant, L. & Ostner, J. (2010). Social bonds enhance reproductive success in male macaques. Current Biology, 20, 2207–2210.

72. Sheldon, B.C., Kruuk, L.E.B. & Alberts, S.C. (2022). The expanding value of long-term studies of individuals in the wild. Nat Ecol Evol, 6, 1799–1801.

73. Shizuka, D. & Johnson, A.E. (2019). How demographic processes shape animal social networks. Behavioral Ecology, 1–11.

74. Silk, J.B., Beehner, J.C., Bergman, T.J., Crockford, C., Engh, A.L., Moscovice, L.R., et al. (2010). Strong and consistent social bonds enhance the longevity of female baboons. Current Biology, 20.

75. Siracusa, E., Boutin, S., Dantzer, B., Lane, J.E., Coltman, D.W. & McAdam, A.G. (2020). Familiar neighbors, but not relatives, enhance fitness in a territorial mammal. Current Biology, 31, 1–8.

76. Snyder-Mackler, N., Burger, J.R., Gaydosh, L., Belsky, D.W., Noppert, G.A., Campos, F.A., et al. (2020). Social determinants of health and survival in humans and other animals. Science (1979), 368.

77. Strauss, E.D. & Holekamp, K.E. (2019). Social alliances improve rank and fitness in convention-based societies. Proc Natl Acad Sci U S A.

78. Temeles, E.J. (1994). The role of neighbours in territorial systems: when are they “dear enemies”? Anim Behav, 47, 339–350.

79. Turner, J.W., Robitaille, A.L., Bills, P.S. & Holekamp, K.E. (2020). Early-life relationships matter: Social position during early life predicts fitness among female spotted hyenas. Journal of Animal Ecology, 1–14.

80. Visser, M.E., Holleman, L.J.M. & Gienapp, P. (2006). Shifts in caterpillar biomass phenology due to climate change and its impact on the breeding biology of an insectivorous bird. Oecologia, 147.

81. Visser, M.E., Van Noordwijk, A.J., Tinbergen, J.M. & Lessells, C.M. (1998). Warmer springs lead to mistimed reproduction in great tits (Parus major). Proceedings of the Royal Society B: Biological Sciences, 265.

82. Ward, A. & Webster, M. (2016). Sociality: The behaviour of group-living animals. Sociality: The Behaviour of Group-Living Animals.

83. Webber, Q.M.R., Albery, G.F., Farine, D.R. & Pinter-wollman, N. (2023). Behavioural ecology at the spatial-social interface. Biological Reviews, 1–40.

84. Wilkin, T.A., Garant, D., Gosler, A.G. & Sheldon, B.C. (2006). Density effects on life-history traits in a wild population of the great tit Parus major: Analyses of long-term data with GIS techniques. Journal of Animal Ecology, 75.

85. Wilkin, T.A., Perrins, C.M. & Sheldon, B.C. (2007). The use of GIS in estimating spatial variation in habitat quality: A case study of lay-date in the Great Tit Parus major. In: *Ibis*.

86. Young, C., Majolo, B., Heistermann, M., Schülke, O. & Ostner, J. (2014). Responses to social and environmental stress are attenuated by strong male bonds in wild macaques. Proc Natl Acad Sci U S A, 111.

